# Iterative Prompt Refinement for Mining Gene Relationships from ChatGPT

**DOI:** 10.1101/2023.12.23.573201

**Authors:** Yibo Chen, Jeffrey Gao, Marius Petruc, Richard D. Hammer, Mihail Popescu, Dong Xu

## Abstract

ChatGPT has demonstrated its potential as a surrogate knowledge graph. Trained on extensive data sources, including open-access publications, peer-reviewed research articles and biomedical websites, ChatGPT extracted information on gene relationships and biological pathways. However, a major challenge is model hallucination, i.e., high false positive rates. To assess and address this challenge, we systematically evaluated ChatGPT’s capacity for predicting gene relationships using GPT-3.5-turbo and GPT-4. Benchmarking against the KEGG Pathway Database as the ground truth, we experimented with diverse prompting strategies, targeting gene relationships of activation, inhibition, and phosphorylation. We introduced an innovative iterative prompt refinement technique. By assessing prompt efficacy using metrics like F-1 score, precision, and recall, GPT-4 was re-engaged to suggest improved prompts. A refined prompt, which combines a specialized role with explanatory text, significantly enhances the performance. Going beyond pairwise gene relationships, we also deciphered complex gene interplays, such as gene interaction chains and pathways pertinent to diseases like non-small cell lung cancer. Direct prompts showed limited success, but “least-to-most” prompting exhibited significant potentials for such network constructions. The methods in this study may be used for some other bioinformatics prediction problems.

## 1. Introduction

Extracting meaningful knowledge from the biomedical literature is becoming increasingly important as the amount of the literature is getting harder and harder to track. One focal point of this endeavor is retrieving information about gene relationships and biological pathways, which offer insights into mechanisms of biological processes and complex diseases like cancer[1]. Historically, computational extraction of these pathways from the biomedical literature has largely depended on rule-based text-mining systems, such as GeneWays[2], Petri net[3], and BioJAKE[4]. While effective to a degree, these methods often confront challenges in accommodating the dynamism and nuance of textual expressions, leading to suboptimal precision and recall. Machine learning methods can replace rule engineering but necessitate vast annotated datasets, making them resource intensive[5]. Some strategies, like distant supervision, have attempted to bypass this limitation, showing potential advantages over traditional rule-based methods[6]. Nonetheless, despite the evolving methodologies, full automation and high accuracy remain elusive due to complexities in literature expression and the intricacies of biological context[7]. Recent studies incorporate images in the biomedical literature using deep learning methods for pathway extractions. However, the performance of this method still suffers from insufficient training data[8]. Large language models, such as ChatGPT, have demonstrated great potentials in pathway extraction[9]^-^ [10], but they are not without challenges; ‘hallucinations’ or instances where these models fabricate non-factual content have posed significant concerns[11]^-^[12].

This study aims to leverage the capabilities of ChatGPT for mining gene relations, a core aspect of pathway extraction. We introduce an innovative iterative prompting methodology and benchmark it on the well-known KEGG Pathway Database[13]. Traditional prompt refinement relies on an evolutionary algorithm that might take extensive computational power[14]^-^[15] or multiple large language models as agents that are hard to adapt[16]^-^[17]. In contrast, our approach fine-tunes prompts and integrates advanced techniques like role prompting, few-shot prompting, and chain-of-thoughts prompting[18], which align the AI-generated content more closely with biological knowledge. Our approach not only addresses the hallucination challenges of ChatGPT, but also seeks to enhance the efficacy and efficiency of pathway extraction.

## 2. Methods

This research employed a hybrid approach combining computational biology and iterative AI prompting to analyze gene interaction triplets within the KEGG Pathway database. A dataset of 300 gene triplets, representative of three fundamental gene interaction types—activation, inhibition, and phosphorylation—was curated to ensure a comprehensive scope. This dataset was divided into subsets for training (200 triplets), validation (50 triplets), and testing (50 triplets), to eliminate assessment biases. The initial set of null prompts was enhanced with role and few-shot prompts, significantly improving result accuracy. Further refinements were achieved through iterative prompt engineering, leveraging GPT-4’s advanced capabilities to incorporate chain-of-thought strategies, resulting in optimal performance after several iterations. In parallel, complex gene relationships were explored, with techniques like chain-of-thought and least-to-most prompting, demonstrating improved coverage of the KEGG Pathway maps.

### 2.1 Data Collection

The KEGG Pathway Database serves as our primary source of the ground truth. KEGG Markup Language (KGML) was used to extract and integrate KEGG Pathway information. A total of 300 gene triplets were randomly selected across different pathway maps with various relations, as shown in Table 1. The triplets contained three major gene interaction types, including activation, inhibition, and phosphorylation, ensuring a comprehensive coverage for our experimental setup.

**Table 1.**
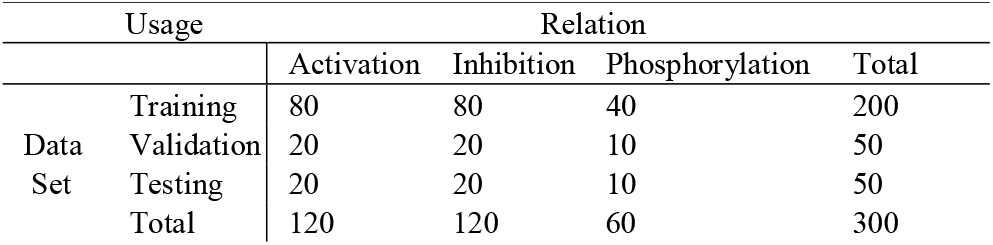
Distribution of gene triplets interactions.

### 2.2 Base Prompt Design

We used various prompt settings, including Role[19]^-^[20] and Few-shot Prompting[21]^-^[22], to explore more accurate answers^16^, as shown below:

#### Null Prompts

- *Prompt Content*: “What effect does {geneA} have on {geneB}?”
- *Format Instruction***: “**Make sure your answer is definitive, composed of ‘activation’, ‘inhibition’, ‘phosphorylation’ or ‘no information’ without further details or explanation.”

#### Designed Prompts with Role, Few-Shots, etc

- *Role Prompting*: “As a computational biologist, determine what is the relation between {geneA} and {geneB} in KEGG Pathway Database.”
- *Knowledge Prompting*: “KEGG pathways are collections of pathway maps that represent our knowledge of the molecular interaction, reaction, and relation networks for various biological processes.”
- *Format Instruction*: “Make sure your answer is definitive, composed of ‘activation’, ‘inhibition’, ‘phosphorylation’ or ‘no information’ without further details or explanation.”
- *Directional Stimulus Prompting*: Emphasizes the importance of directionality in the relationship.
- *Few-Shot Prompting*: Q: What effect does gene EGF have on gene EGFR? A: Activation. Q: What effect does gene GRK2 have on gene OR2AJ1? A: Inhibition. Q: What effect does gene CDK9 have on gene NELFB? A: Phosphorylation. Q: What effect does “{gene1}” have on “{gene2}” directly?

### 2.3 Iterative Refinement with GPT-4[23]^-^[24]

To further enhance the performance, we leveraged GPT-4’s capabilities for iterative prompt refinement[25]^-^[26]. The model was guided to refine its answers through techniques like role prompting, omitting some details, and incorporating chain-of-thought or tree-of-thoughts[27] strategies.

#### 2.3.1 Prompts for mining gene relations

In mining gene relations, we employed prompts analogous to those delineated in Section 2.2. When interfacing with the OpenAI API, a prompt is divided into two principal components: the system content and the user content. As delineated in OpenAI’s official documentation, the system content calibrates the assistant’s behavior, while the user content conveys specific queries or comments, eliciting responses from the assistant. In our method, the bulk of the prompt is integrated into the system content, reserving the actual query for the user content.

#### 2.3.2 Meta-Prompt Design

As illustrated in Figure 1, the meta-prompt functions as a guiding schema for GPT-4, steering it towards iterative refinement of prompts based on their prior performance. The exemplar meta-prompt employed in our study is as follows:

**Fig. 1.**
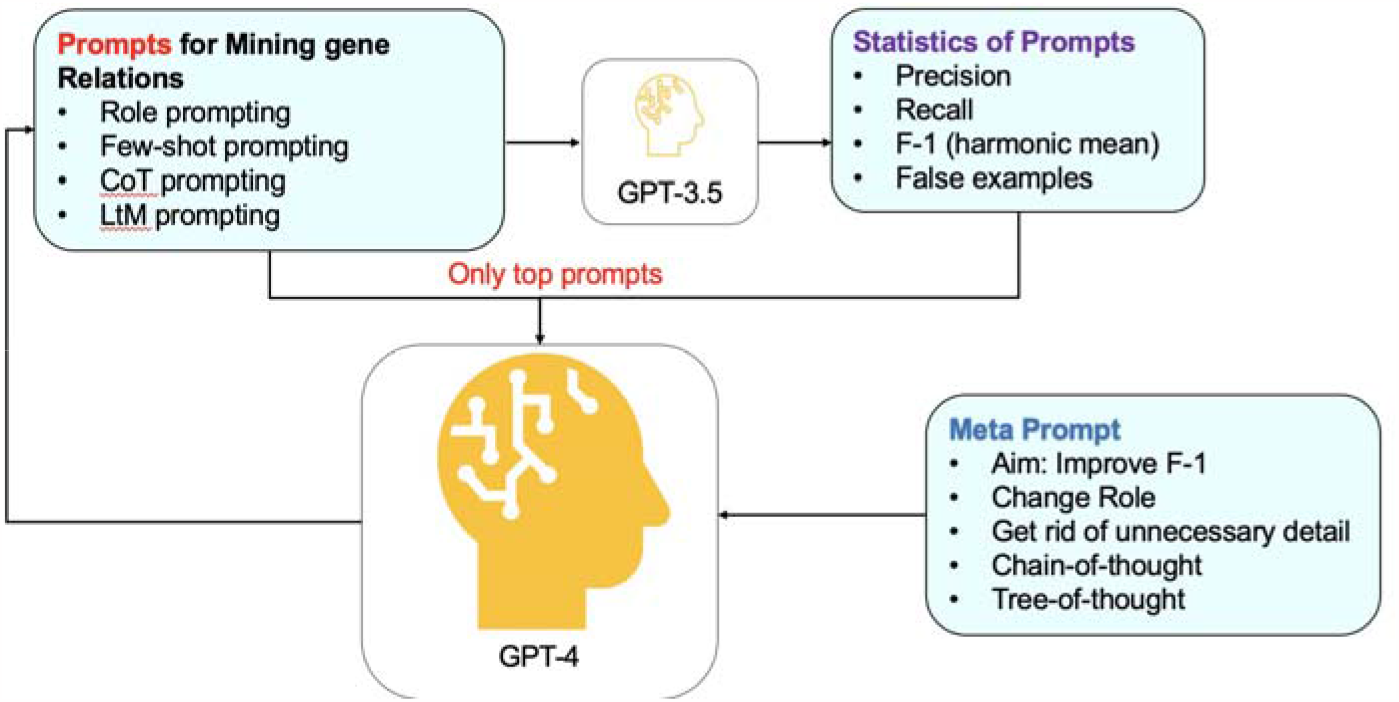
Iterative Prompt Refinement Framework Utilizing GPT Models for Gene Relation Mining. The framework initiates with diverse prompting strategies—role-based, few-shot, chain-of-thought, and least-to-most prompting—to extract gene relations. These prompts undergo an initial evaluation by GPT-3.5, selected for its efficiency in processing factual queries. The prompts’ effectiveness is quantified using precision, recall, and F-1 scores, alongside the identification of false positives. Top-performing prompts, alongside a meta-prompt, inform the subsequent iteration with GPT-4, which refines the prompts further. The meta-prompt outlines the objective of enhancing the F-1 score, directs the model to modify prompt roles, eliminate extra details, and apply advanced reasoning frameworks such as chain-of-thought and tree-of-thought. This approach exploits GPT-4’s advanced logical capabilities to autonomously improve prompts, leveraging the lower cost and higher speed of GPT-3.5 for initial fact-based query processing.

*“As a senior professional prompt engineer, refine former prompts focused on direct gene relations, including activation, inhibition and phosphorylation in the KEGG Pathway database. The goal is to generate a prompt that improves the accuracy when determining the effect one gene has on another, with the correct answer corresponding to the KEGG Pathway database. Ensure that the new prompt is original, has never appeared before, and that it either enhances the existing prompts’ accuracy or changes the content of prompts dramatically. The prompt should be applicable to a broad range of genes without focusing on specific genes, gene families, or particular aspects of gene phosphorylation relations. The response should be definitive, selected from “activation”, “inhibition”, or “phosphorylation” and only composed of these words separated by “*…*” without further details*.

*You will be provided with some top-performed prompts and their statistics, on integrating the following top-performed prompts, please think about how to integrate these prompts in a step-by-step manner. You can try to answer the following question before providing the revised prompt: 1. What are the key elements from each prompt that contribute to their respective performance scores (overall_F-1, activation_F-1, inhibition_F-1, phosphorylation_F-1)? 2. Are there any specific terms or phrases in the prompts that could be refined or replaced to make the instructions clearer for the model? 3. Are there any redundant or repetitive phrases that can be removed or combined to create a more concise and effective prompt? 4. How can the context and background information be integrated without making the prompt too long or complex, while still providing the model with enough information to answer accurately? 5. Is there any relevant information or context that is missing from the original prompts that could be added to improve the model’s understanding and accuracy?*

*Below are detailed instructions on how to generate better system content, user question and temperature: 1. Please change the role of the original prompt such as “Act as a bioinformatic researcher”, but make sure you are changing to another relevant but original role that has never appeared before. 2. Add details to the instruction. 3. Get rid of unnecessary detail. 4.Please make sure your refined prompt includes few-shot examples of gene relations but you may change the examples. 5. Taking all the previous example prompts and their accuracy into consideration, you might want to integrate the prompts with the best performance. 6. If you believe that current prompts cannot be improved, you can change the original prompt dramatically. 7. The response should include the revised system content and user question but in a python code format. 8. Change the temperature to a temperature that you think will help GPT to respond more accurately. In general, higher temperature makes the question more diverse and unpredictable while low temperature makes it more conservative. 9. The response should include the answers to the step-by-step questions, revised system content, user question and temperature, separated by ‘---’. Your answer should be structured like “answers:* … *--- system content:* … *--- user question:* … *--- temperature:* …*”. 10. Make sure your revised question includes “{gene1}” and “{gene2}” to indicate the starter gene and receiver gene*.

*Please answer these questions before providing the revised prompt, user question and temperature separated by ‘---’*.

*Do not provide your answer to these questions in your response.*

*For your information, this is the previous system content:”*

The design rationale of the meta-prompt is anchored in a multi-faceted approach:

1. Role Conditioning: By configuring GPT-4’s perspective to that of a “prompt engineer,” we anticipated fostering a heightened quality of response. This role-play encourages a more precise and methodological approach to prompt refinement.
2. Task Goal Communication: Explicitly defining the purpose of the prompt—to mine gene relations—and outlining the expected response structure provides the model with clear context and expectations.
3. Guided Thought Process: By introducing a series of interrogative directives, we prompted GPT-4 to critically assess and understand the nuances of effective prompt design, thereby fostering a more considered response.
4. Detailed Instructional Guidance: Comprehensive directives, ranging from modifications to existing roles to details to be included or excluded, ensure the model has a structured pathway to elicit improvements. For instance, emphasizing the need for originality in roles while still staying pertinent, steers the model away from redundant or overused contexts. Moreover, including performance statistics from previous prompts provides GPT-4 with empirical data, potentially guiding its refinement process based on proven efficacy.

This design aims to optimize the trade-off between prompt brevity and informativeness, ensuring the meta-prompt serves as an effective tool for eliciting high-quality refinements from GPT-4.

#### 2.3.3 Hyper-parameters configuration

In our prompt-refinement algorithm, two pivotal parameters govern the outcome of prompt engineering: the temperature and the number of prompts. We implemented a linear temperature adjustment scheme. Specifically, for a given iteration *i* with a starting temperature *S* and an ending temperature *E*, the temperature for the *i*^th^ iteration, *T*_i_, is computed in Eq. (1):

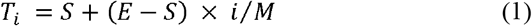

where *M* represents the total number of iterations. Moreover, the parameter “number of prompts” determines the number of top prompts GPT-4 is exposed to before offering revised prompts. For example, when the number of prompts equals 2, GPT-4 will be provided with top 2 previous prompts.

For hyper-parameter optimization, we employed a grid search strategy. Starting and ending temperatures were selected from the set {0, 0.25, 0.5, 0.75, 1}, while the potential values for the number of prompts were {1, 2, 3}. Each configuration was iterated ten times under consistent meta-prompt and other parameters. We utilized the best-performing prompts from these iterations on a validation set, with performance gauged by the validation error. Our empirical findings highlighted optimal settings: a starting temperature of 0.5, an ending temperature of 0.75 (Appendix 1). Statistical analyses revealed that the discrepancy between the starting and ending temperatures, as well as the average temperature across iterations, significantly correlated with the validation error.

### 2.4 Iterative Prompt Refinement with Tree-of-Thought Framework

Our study employed the “Tree-of-Thought” framework, an innovative approach developed by Yao et el.[27] and Long[28] to emulate the human mind’s method of tackling complex reasoning tasks. This framework systematically explores solution spaces, emulating a structured, tree-like thought process akin to human cognitive strategies. A distinctive feature of this framework is its capacity for strategic backtracking, which is crucial in overcoming impasses and enhancing problem-solving efficiency.

A key aspect of the tree-of-thought framework is its adaptability, achieved through its compatibility with various language models that form its operational backbone. This adaptability is essential for addressing various computational and linguistic challenges, extending the framework’s applicability across numerous fields. Further augmenting its robustness, the framework incorporates a suite of search strategies, including depth-first search, breadth-first search, and Monte Carlo search[29]. This multifaceted approach not only supports the framework’s resilience in diverse task environments but also amplifies its overall analytical effectiveness.

Our project utilized the tree-of-thought framework in conjunction with the GPT-4 language model, leveraging its advanced capabilities in language processing. This integration was critical for enhancing the framework’s analytical and problem-solving capacities. A tree-of-thought framework implemented by Agora (accessible at https://github.com/kyegomez/tree-of-thoughts), based on the Monte Carlo search algorithm, was used in our study. We configured the framework to generate a single thought per iteration, allowing up to three steps and four stages with 0.5 as tree-pruning threshold in the decision-making process before deriving a conclusion to facilitate iterative refinement of prompts in our interactions with GPT-4.

## 3. Results

The objective of this study was multifaceted: to assess variabilities in model performance, determine the potential advantages of designed prompts, and understand the viability of iterative refinement strategies. Our findings revealed stark temporal fluctuations in GPT-3.5-turbo’s performance, with a significant F-1 score reduction over a few months. Notwithstanding these temporal variations, the employment of role-based and few-shot prompts demonstrated appreciable enhancements in performance, particularly as the model aged. Furthermore, this approach showcased GPT-4 model’s self-reflection and self-improvement capabilities.

### 3.1. Initial Testing Performance

To ascertain the efficacy of our methodology, we first analyzed the temporal variability of the GPT-3.5-turbo model (Fig. 2). In March 2023, querying the model regarding gene interrelationships yielded an F-1 score of 0.504. Notably, by July 2023, this score dramatically decreased to 0.068, confirming a prior study that suggests significant temporal variability in GPT’s performance[30]. Yet, as our results show, the advantage of careful prompt design, utilizing role-based and few-shot approaches, becomes increasingly pronounced with time. Specifically, we observed improvements from 3% by using role and few-shot prompting in March, 80% in May, and 181% in July, respectively.

**Fig. 2.**
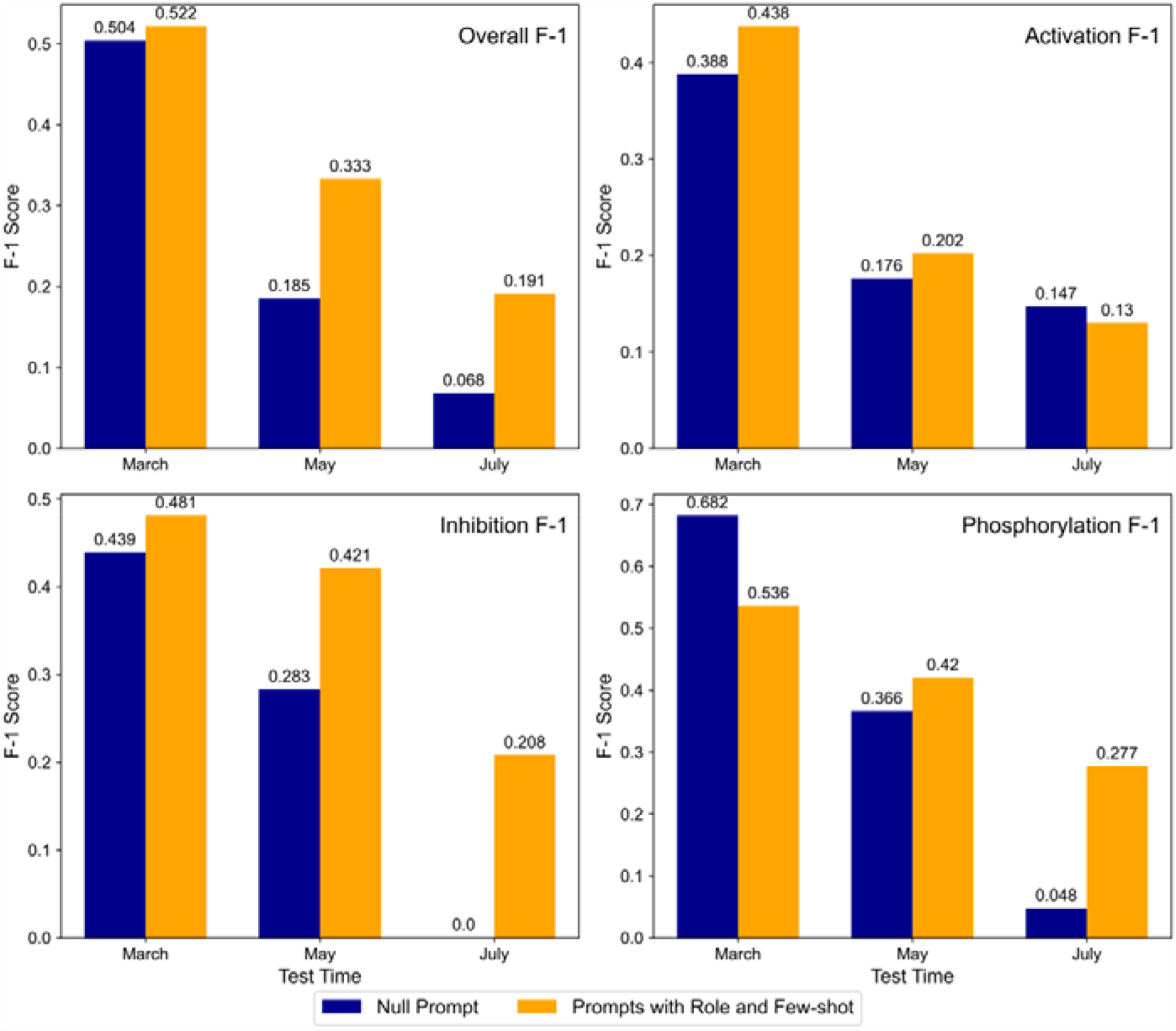
Temporal variability of F-1 scores for gene interaction. Each bar chart in the figure presents the F-1 performance metric for a specific type of gene interaction—phosphorylation, activation, inhibition, and overall interaction—assessed at three distinct time points: March, May, and July 2023. The null prompt approach (blue) establishes a baseline, while the few-shot prompt method (orange) represents an advanced strategy incorporating role-based, few-shot learning. Contrary to the expectation, the performance scores, particularly for the overall, inhibition, and phosphorylation interactions, decreased over the assessed months.

### 3.2. Iterative Refinement

Transformers, like GPT-3.5, inherently produce predictions in a sequential manner, devoid of self-reflection on prior outputs. Harnessing this characteristic, we devised an iterative refinement protocol. In this scheme, GPT-3.5 is initially tasked with generating gene relation prompts. Subsequent evaluation against the KEGG pathway database provides an initial performance metric. Drawing upon these outputs, GPT-4 refined prompts to bolster the F-1 score. This refinement employs strategies including role prompting, chain-of-thought, and tree-of-thoughts. The results of this iterative approach, albeit promising initially, displayed variability in subsequent runs.

**Fig. 3.**
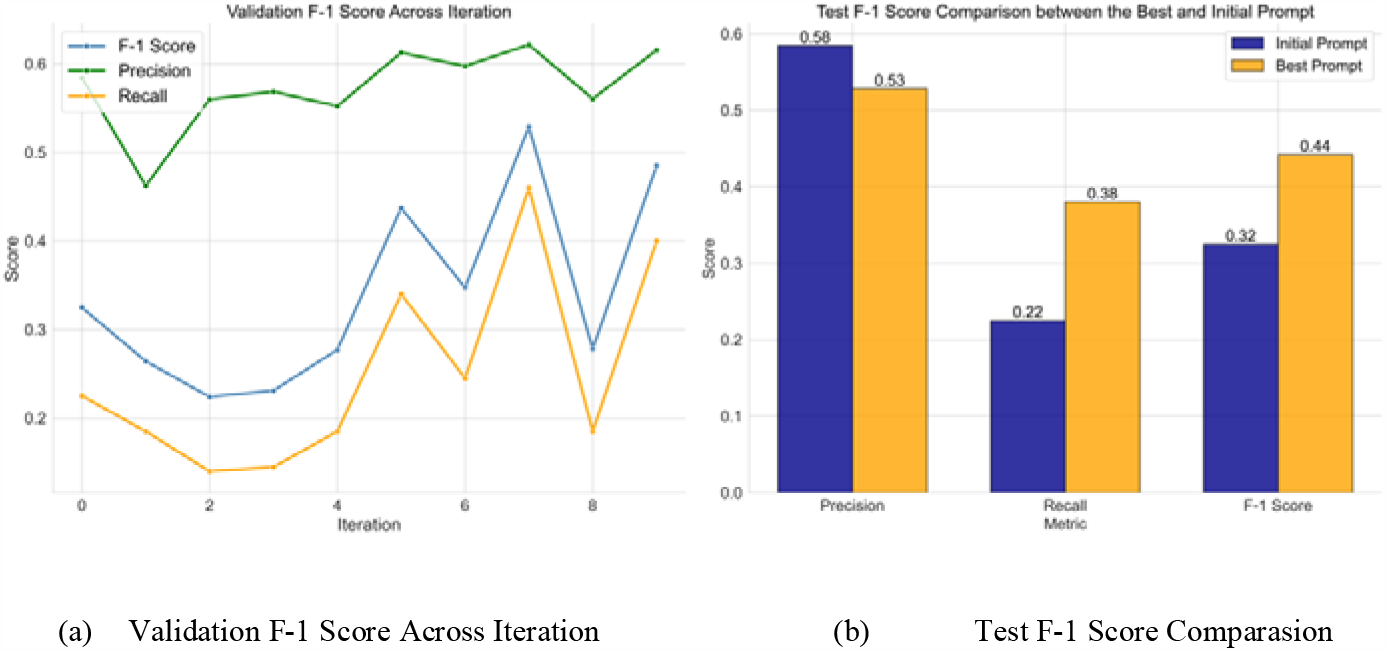
Iterative refinement and comparative analysis of performance metrics. Subfigure (a) delineates the trajectory of precision, recall, and F-1 scores throughout the iterative refinement process. Each line represents the progression of a specific metric, with annotated points indicating the maximum and initial values achieved. This visualization highlights the algorithm’s dynamic adjustment and the evolution of model performance over iterations. Subfigure (b) is a bar plot that contrasts the initial and optimally refined prompts, illustrating significant performance enhancements. The comparison underscores the improvement in precision, recall, and F-1 scores, demonstrating the iterative refinement’s effectiveness in optimizing prompts.

Below is the prompt revised by ChatGPT:

*“****As a molecular biology specializing in gene interactions***, *determine the direct relationship between {gene1} and {gene2} in the KEGG Pathway Database, which represents molecular interactions, reactions, and relation networks in biological processes. Provide a definitive answer using one or more of the following terms: “activation”* ***(gene1 activates gene2)***, *“inhibition”* ***(gene1 inhibits gene2)***, *“phosphorylation”* ***(gene1 phosphorylates gene2)***, *or “unknown”* ***(interaction is uncertain or not present)***. *Separate multiple terms with “*…*”. Your answer should only include the relationship terms without additional context. For example:*

*Q: What is the direct effect of gene EGF on gene EGFR? A: Activation*.

*Q: What is the direct effect of gene GRK2 on gene OR2AJ1? A: Inhibition*.

*Q: What is the direct effect of gene CDK9 on gene NELFB? A: Phosphorylation*.

*Q: What is the direct effect of gene “{gene1}” on gene “{gene2}”? A:”*

The selected prompt from ChatGPT notably modified the role to “a molecular biologist specializing in gene interactions” and provided concise explanations for each relation. This modification resulted in a significant improvement in the outcomes. This iterative process culminated in an F-1 score of 0.439 on the validation set, closely mirrored by an F-1 of 0.441 on the testing set.

### 3.3. Iterative Prompt Refinement with the Tree-of-Thoughts Framework

The tree-of-thoughts framework with GPT-4 as the backbone generates improved versions of prompts (system contents) and questions previously fed to GPT-3.5. With a detailed meta-prompt giving commands and suggestions, the framework improved prompt quality. Another variable of temperature also affected the accuracy and consistency of the system contents and questions. These new contents were again fed to GPT-3.5 to mine gene relationships and form potential knowledge graphs, improving upon the innovative approach of mining with LLMs. By introducing specialized roles and explanatory text, and giving options to remove/add information and context to the contents and questions, the tree-of-thoughts framework improved the accuracy and F-1 score of the gene relationships. Repeated contents and certain roles significantly improved the consistency of GPT-3.5’s response.

The tree-of-thoughts framework accepts two additional parameters: the number of prompts to generate and the range of temperature. More prompts increase variability and decrease accuracy as it becomes harder to choose the best prompt. The range of temperatures has no dramatic effect as the values tend to be 0, but higher temperature ranges increase variability in generated prompts.

#### 3.3.1. Meta-Prompt

The meta-prompt was based on the knowledge gathered from previous experiments: role prompting, generated knowledge prompting, formatting instructions, directional stimulus prompting, and few-shot prompting. Through suggesting role changes, context and background edits, emphasis on directionality and tweaking examples, the meta-prompt instructed the tree-of-thoughts framework to improve GPT-3.5’s response. Formatting instructions for GPT-3.5’s response was to enhance response parsing and accuracy, which led to a trend in lower temperature (see Section 3.3.4). The initial meta-prompt (Appendix 2) had three main experiments conducted: varying language, organizing structure, and editing steps of generating new prompts. The switch from requests to commands (i.e., “do <x>” to replace “please try <x>”) had no explicit effect. Both the framework and GPT-3.5 have flexibility in interpreting contents, with other variables affecting accuracy, precision, and recall more heavily.

The structure of the meta-prompt is important for consistency and accuracy. The most effective organization was also the clearest: three sections of background and context, instructions for content and questions, and suggested changes. This increased the F-1 score from an average of 0.10 to 0.18. This reduced redundancies while maintaining and even improving accuracy and the F-1 score given that the other parameters were constant.

The final meta-prompt is shown in Appendix 3. Adding new instructions, such as to shorten the system content as much as possible, or “to change the number of examples of gene relations”, saw varying results. “Changing the number of examples” and other similar modifications saw a minimal effect on the average F-1 score (0.01 increase). More importantly, the F-1 scores across the three types of relationships changed in both positive and negative directions, falling within GPT-3.5’s bounds of inconsistency (See Section 3.3.2 for the effects of shorter system contents).

Explicit instructions to modify the prompt in a certain way (such as changing the role to a “molecular geneticist”) had no significant effect (Appendix Table 1), except when the tree-of-thoughts was commanded to “implement the answers to <rhetorical questions> when changing the old prompts”. This improved the F-1 score from 0.20 to 0.32 (Appendix Table 2). Other explicit instructions such as “avoid false positives” and “reduce errors” were found to improve recall in the F-1 score.

#### 3.3.2. System Content

The system content, or the past prompts fed into the tree-of-thoughts, were a low factor of change in accuracy and F-1 score unless they were repeated. More examples of system content did not change the revised contents returned by the tree-of-thoughts, as all contents were similar in length and content with slight changes in roles and examples for GPT-3.5. However, the many contents did increase the F-1 score of the first iteration to an average of 0.30 (Appendix Table 2). Providing 5 contents returned an F-1 score of 0.35 while only one content resulted in an F-1 score of 0.10 average over 20 test cases. This pattern can be attributed to GPT-3.5’s processing of system content, where it may heavily favor repeated language over others. Its greatest effect was decreasing the number of responses with “no information”, improving consistency and F-1 score.

Meta-prompt changes to generate content of “minimal length” resulted in content with three sentences: one providing a role and requesting the GPT-3.5 to determine the gene relationship (role and generated knowledge prompting); one giving the four possible responses (format instructions); and one reinforcing that the response can only be one of these four options (format instructions). Directional stimulus and few-shot prompting showed up less often in the shorter contents and did not have a great impact on GPT-3.5’s response.

Weak system contents only requesting gene pathways and providing no extra information were still revised to the quality of the three-sentence examples. These improved contents resulted in F-1 scores increasing from 0.03 to 0.14, which reinforces that the framework works exceptionally and that GPT-3.5 reports more accurate gene relationships with better content.

#### 3.3.3. Effect of Compression on Performance

GPT-3.5 and many other LLMs count processing power by tokens, which restricts iterations, processing, and flexibility with experimentation. However, through human editing of the meta-prompt, the tree-of-thoughts framework can compress all queries to use fewer tokens. The meta-prompt itself can be fed through the tree-of-thoughts framework, but was edited manually for these trials. The meta-prompt was requested to compress the content to three or one sentences, which it achieved in all trials without shortcuts such as run-on sentences. Even with the decrease in quantity, the response of GPT-3.5 to the optimized content and question equaled or outperformed the accuracy and F-1 score of the longer versions. Despite the directional stimulus and few-shot prompting in contents (see Section 3.3.2) or generated knowledge prompting and formatting instructions in questions, GPT-3.5 still provided parsable responses that promised results. The detriment of using fewer tokens is combated by the natural optimization of the tree-of-thoughts framework.

### 3.4. Complex Gene Interaction Mining

Our exploration did not stop at direct gene relations. Venturing into complex interactions, an interaction chain between EGFR and ERK was studied. Impressively, ChatGPT’s findings mirrored the KEGG Pathway map on melanoma, reinforcing the notion that in-depth gene relationships could be mined from language models. Details on the prompts and responses of ChatGPT can be found in Appendix 4.

We further reconstructed the comprehensive gene pathway for non-small cell lung cancer. Our initial approach involved a direct inquiry into ChatGPT to reconstruct this gene pathway. The visual representation in Neo4j depicted only 7 genes, devoid of any gene relationships. Of these, 6 genes aligned with the original KEGG pathway map, encompassing 19.3% of the entirety of the KEGG Pathway map (36 genes in total, Appendix 5). Acknowledging the shortcomings of these results, we explored an optimization strategy. An initial consideration was the chain-of-thought method, which did not yield the desired efficacy in this context. Consequently, we turned to the ‘Least to Most’ (LtM) prompting technique.

The “Least-to-Most” (LtM) prompting technique stands as an evolutionary advancement over the traditional chain-of-thought prompting. Where chain-of-thought has showcased remarkable proficiency in numerous natural language reasoning tasks, its efficacy diminishes when confronted with complex problems surpassing the scope of its initial prompts. This recognized limitation spurred the development of the LtM strategy. At its core, the LtM approach systematically deconstructs intricate problems into an ordered sequence of more manageable subproblems. Each subproblem’s resolution not only paves the way for addressing subsequent challenges but also enhances the precision of subsequent prompts by integrating insights from previously solved subproblems. Empirical data reinforces the prowess of this methodology, especially when combined with chain-ofthought.

We deployed this strategy by first asking ChatGPT what subquestions could be solved before it reconstructed the pathway (Appendix 6). Then we ask each question sequentially with previous questions and answers provided through context prompting. Finally, when we threw out our final question, where we asked ChatGPT to reconstruct the gene pathway about non-small cell lung cancer, we were able to identify 16 genes, comprising 21 activation and 1 phosphorylation relations. Of these, 10 genes correlated with the original KEGG pathway map, covering approximately 27.8%. While these findings represent a tangible advancement compared to the initial results, there remains substantial room for enhancement.

**Fig. 4.**
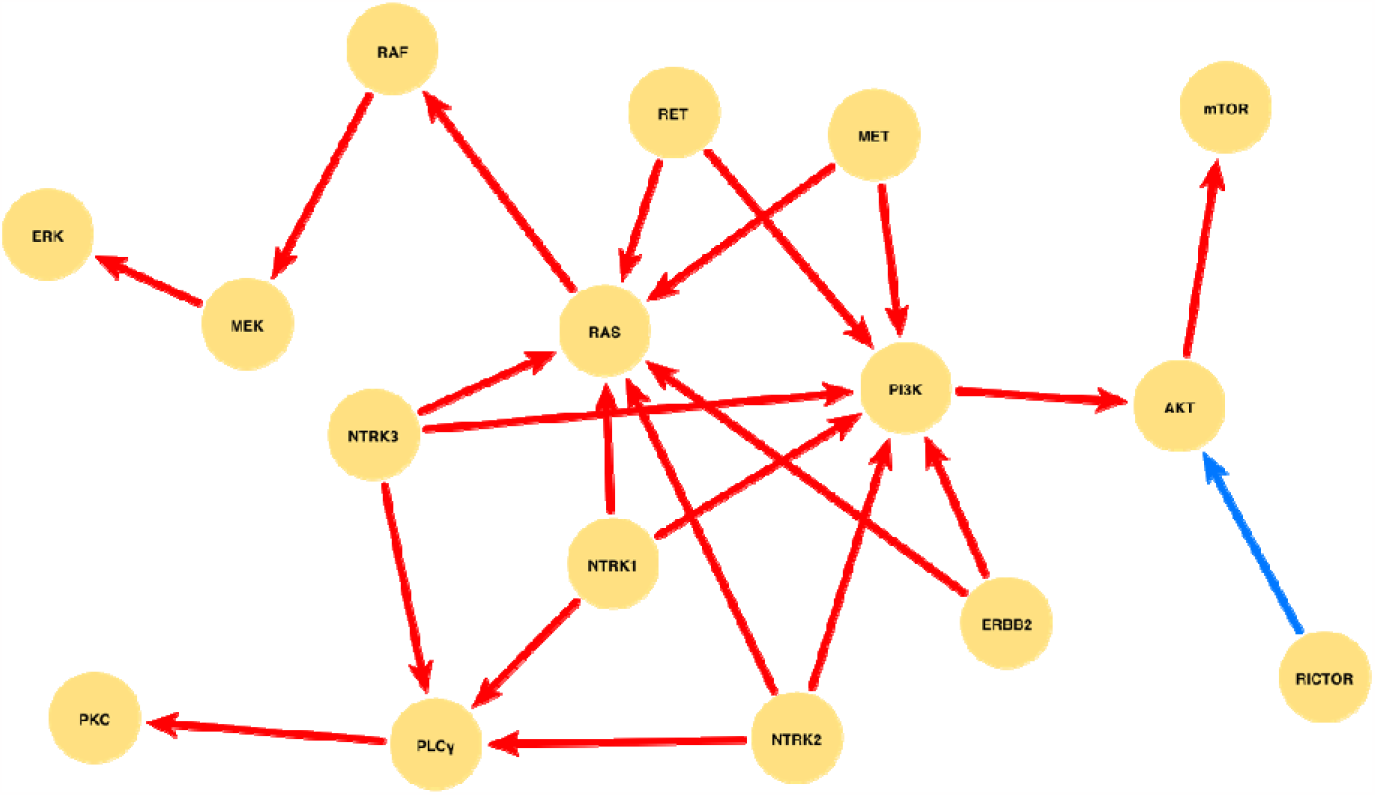
Enhanced visualization of the non-small cell lung cancer gene pathway post-implementation of the ‘Least-to-Most’ (LtM) prompting technique. This advanced representation identifies 16 interconnected genes, encompassing 21 activation (depicted in red) and 1 phosphorylation (depicted in blue) relations. Approximately 27.8% of the genes (highlighted in yellow) correlate seamlessly with the original KEGG pathway map, underscoring the enhanced depth and accuracy attained through the LtM approach.

## 4. Discussion and Conclusion

This study introduces a methodological approach to harness gene relationships from large language models. Leveraging the KEGG Pathway Database as a benchmark, our dataset consisted of 200 training gene triplets, 50 validation sets, and an additional 50 sets for testing, spanning three primary gene interaction types. Our tests were conducted on the GPT-3.5-turbo version available via OpenAI’s API. Further, we incorporated role prompting and few-shot prompting, emphasizing the directionality of relationships. Our results showcased a marked performance enhancement with this tactic, relative to the null prompts. A significant premise for this design was the conditioning ability of role prompting on models like ChatGPT. A pivotal observation was the pronounced increase in the F-1 score, albeit with room for further enhancement.

Recognizing the directional prediction nature of transformers, we hypothesized the efficacy of prompting refinement through iterative means. Our strategy involved deploying the mined gene relations into GPT-3.5 and comparing outputs with the KEGG pathway database, focusing on precision, recall, F-1 scores, and highlighting inaccuracies. Feedback was then provided to GPT-4 to refine the F-1 score, utilizing techniques like role prompting, chain-of-thought, or tree-of-thoughts. Furthermore, we explored complex gene relationships by extending our scope beyond direct gene connections. One such endeavor focused on deciphering the interaction chain between EGFR and ERK. ChatGPT’s findings echoed the KEGG Pathway map on melanoma, delineating the activation chain from EGFR to ERK.

Finally, we reconstructed the entire gene pathway for non-small cell lung cancer using GPT. Initial attempts yielded suboptimal results, with only 19.3% coverage of the original KEGG Pathway map. Given the limited success of the chain-of-thought approach, we pivoted to the LtM prompting technique, rooted in educational paradigms. This methodology yielded some improvement, underscoring the complexities and potential avenues for refinement in gene pathway reconstructions.

In conclusion, our findings underscore the potential and challenges of harnessing large language models in bioinformatics. The fluctuating performance metrics, the susceptibility to training data, and the necessity for iterative refinement strategies emphasize the dynamic nature of this research area. As we continue to fine-tune our approaches and understand the intricacies of these models, it opens new doors to novel methods of data extraction and interpretation. Given the vast and diverse training data of models like ChatGPT, the ability to extract specialized knowledge also offers an intriguing avenue for diverse fields, not limited to bioinformatics.

Our investigation into extracting gene relationships from large language models yielded several key findings:

- **Performance Variability**: The GPT-3.5-turbo’s performance in extracting gene
- relationships exhibited notable variability over time. While initial tests showed promising F-1 scores, subsequent evaluations marked a substantial decrease.
- **Prompt Refinement**: The implementation of role prompting and few-shot
- prompting showcased enhanced performance, especially in deciphering the directionality of relationships. This suggests that the strategic structuring of prompts can potentially unlock more accurate information from these models.
- **Iterative Feedback Loop**: An iterative feedback loop with GPT-4 demonstrated peak performance at the third iteration. However, the subsequent fluctuations underscore the inherent challenges and the necessity for consistent refinements.
- **Complex Gene Relationships**: When mining for complex gene relationships, the model’s response resonated well with established pathway maps, suggesting its potential in understanding multifaceted gene relationships. Our method’s success emphasizes the significant potential of mining knowledge from large language models. However, the challenges we faced, particularly the variable performance metrics and the model’s susceptibility to training data nuances, shed light on areas for future work. Potential avenues include:
- **Model Customization**: Exploring other language models specifically open-source models such as Llama 2, for bioinformatics tasks, as they may offer more transparent and specialized outputs.
- **Refinement Algorithms**: Developing more advanced algorithms for iterative prompting, possibly integrating advanced AI techniques to better guide the model towards desired outputs.
- **Broader Applications**: Evaluating the method’s efficacy across more research problems, expanding its utility beyond gene relationships.

## 5. Acknowledgments

This work was supported by an NIH-NLM grant R01-LM013392.

## Appendix 1. Heatmap Analyses of Iterative Refinement Algorithm Grid Search Results

**Appendix Figure 1.**
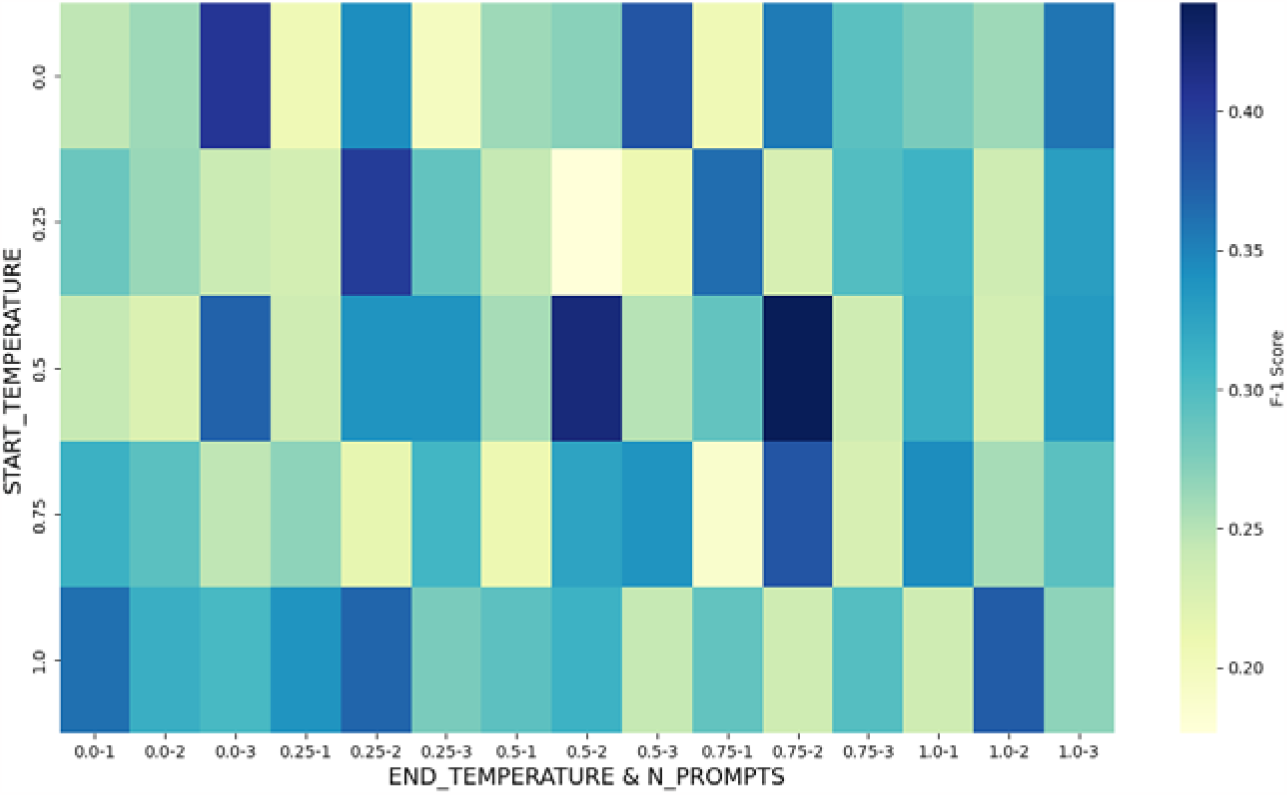
Heatmap representation of grid search results for the iterative refinement algorithm based on F-1 Scores. Different shades highlight the effectiveness of each parameter combination in refining the model’s performance.

**Appendix Figure 2.**
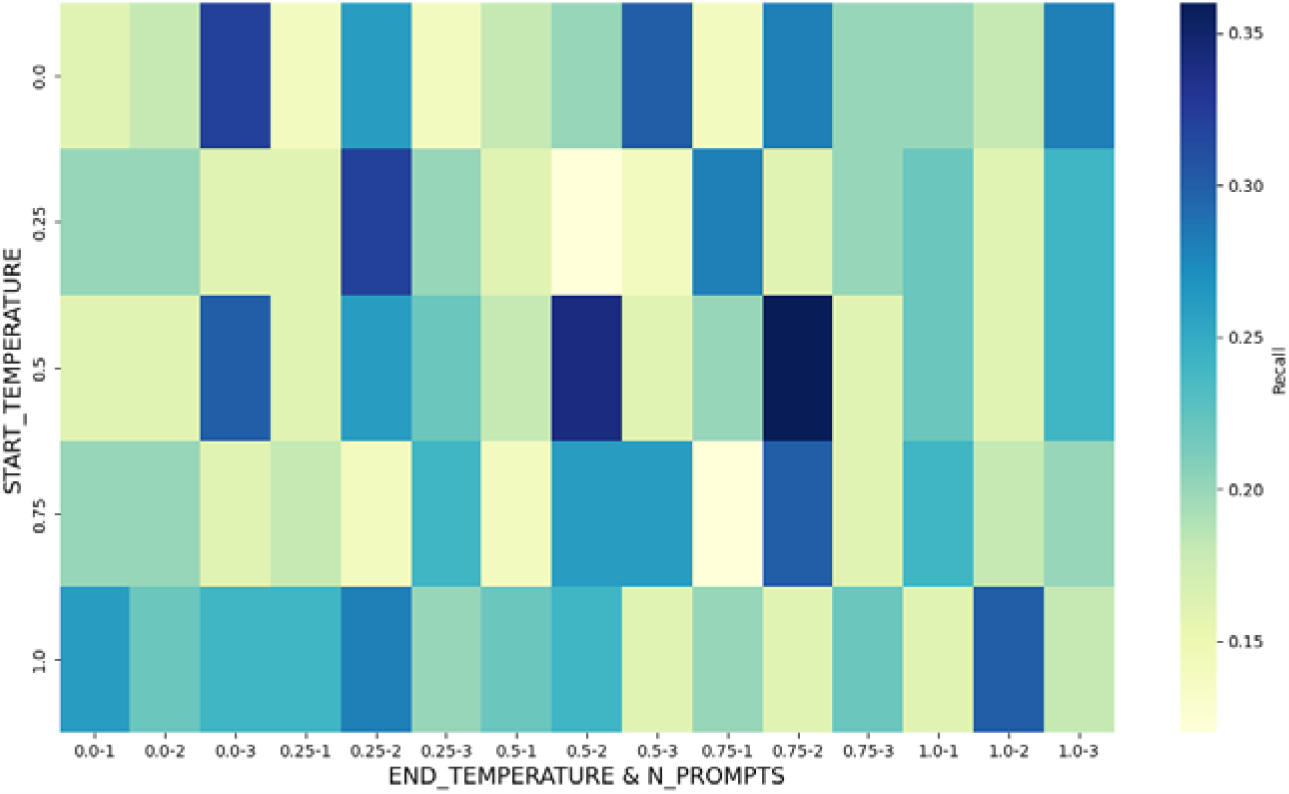
Heatmap visualization displaying grid search results of the iterative refinement algorithm based on Precision metrics. Variations in color intensity underscore the differential impact of each hyperparameter setting.

**Appendix Figure 3.**
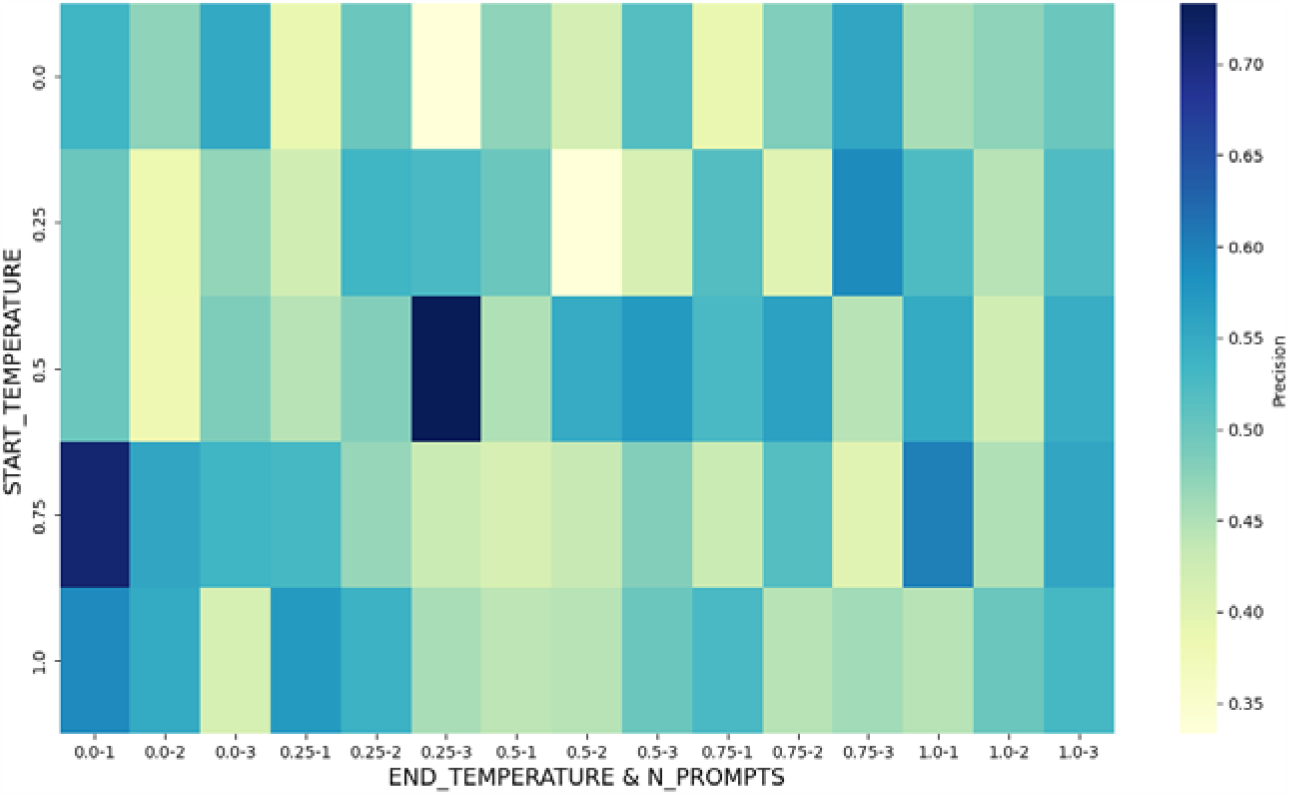
Heatmap plot showcasing grid search results for the iterative refinement algorithm, focusing on Recall metrics. The gradient in color marks the range of outcomes achieved with different parameter combinations.

## Appendix 2: Initial Meta-Prompt

*As a senior professional prompt engineer, refine former prompts focused on direct gene relations, including activation, inhibition and phosphorylation in the KEGG Pathway database. The goal is to generate a prompt that improves the accuracy when determining the effect one gene has on another, with the correct answer corresponding to the KEGG Pathway database. Ensure that the new prompt is original, has never appeared before, and that it either enhances the existing prompts’ accuracy or changes the content of prompts dramatically, the prompt should be applicable to a broad range of genes without focusing on specific genes, gene families, or particular aspects of gene phosphorylation relations. The response should be definitive, selected from “activation”, “inhibition”, or “phosphorylation” and only composed of these words separated by “*…*” without further details*.

*You will be provided with some top-performed prompts and their statistics, on integrating the following top-performed prompts, please think about how to integrate these prompts in a step-by-step manner. You can try to answer the following question before providing the revised prompt: 1. What are the key elements from each prompt that contribute to their respective performance scores (overall_F-1, activation_F-1, inhibition_F-1, phosphorylation_F-1)? 2. Are there any specific terms or phrases in the prompts that could be refined or replaced to make the instructions clearer for the model? 3. Are there any redundant or repetitive phrases that can be removed or combined to create a more concise and effective prompt? 4. How can the context and background information be integrated without making the prompt too long or complex, while still providing the model with enough information to answer accurately? 5. Is there any relevant information or context missing from the original prompts that could be added to improve the model’s understanding and accuracy?*

*Below are detailed instructions on how to generate better system content, user question and temperature: 1. Please change the role of the original prompt such as “Act as a bioinformatic researcher”, but make sure you are changing to another relevant but original role that has never appeared before 2. Add details to the instruction. 3. Get rid of unnecessary detail. 4. Change the example to relations among other examples. But please make sure your refined prompt includes examples. 5. Take all the previous example prompts and their accuracy into consideration; you might want to integrate the prompts with the best performance. 6. If you believe that current prompts cannot be improved, you can change the original prompt dramatically. 7. The response should include the revised system content and user question but in a python code format. 8. Change the temperature to a temperature that you think will help GPT to respond more accurately. In general, higher temperature makes the question more diverse and unpredictable while low temperature makes it more conservative. 9. The response should include the answers to the step-by-step questions, revised system content, user question and temperature, separated by ‘---’. Your answer should be structured like “answers:* … *--*

*- system content:* … *--- user question:* … *--- temperature:* …*” 10. Make sure your revised question includes “{gene1}” and “{gene2}” to indicate the starter gene and receiver gene*.

*Do not provide your answer to these questions in your response.*

*For your information, this is the previous system content:*

## Appendix 3: Final Meta-Prompt

*As a senior professional prompt engineer, refine former prompts focused on the direct gene relations of activation, inhibition and phosphorylation in the Kegg Pathway database. You must generate a prompt to feed to a model that responds with the effect of one given gene on another with maximum accuracy compared to the ground truth, the Kegg database. The new prompt should always be original and either enhance existing accuracy or change contents dramatically. It should be applicable to all genes without focusing on any specific families, relations, etc. and be of minimal length. The returned prompt should always include the revised system content and user question, and temperature in Python code format, separated by ‘---’, and formatted like this:*

*“answers:* … *--- system content:* … *--- user question:* … *--- temperature:* …*”*.

*Ensure the revised question includes “{gene1}” and “{gene2}” to indicate the starter and receiver genes. The model’s response to the prompt should be strictly one or two words from: “activation”, “inhibition”, and “phosphorylation” separated by “*…*”*.

*You will be provided with top-performing prompts. Try to answer the following five questions when improving these prompts:*

*What are the key elements from each prompt that contribute to their accuracy?*

*What specific terms or phrases in the prompts should be edited to clarify instructions to improve accuracy?*

*What redundancies should be edited to create a shorter prompt while still retaining accuracy?*

*What necessary context and background information should be edited to be concise and improve accuracy?*

*What changes can be made to decrease errors (“no information” instead of a correct relation and “phosphorylation” instead of “activation”)?*

*Please answer these questions and use these answers to revise the prompt before providing the new prompt, user question and temperature separated by ‘---’. Do not provide your answers to these questions in your response*.

*Below are six examples to generate better system content, user questions, and temperature:*

*Change the role of the original prompt to another relevant but original role such as “Act as a molecular geneticist”*.

*Add details to the instruction, such as instructing the model to simulate gene pathways to figure out relationships or search the Kegg database*.

*Get rid of unnecessary details, such as the origins of the Kegg database*.

*Change the number of examples or example relations to other 100% true examples*.

*Rephrase the user question to be more specific, such as “What effect does protein {gene1} have on protein {gene2} directly out of the following: activation, inhibition, phosphorylation, or no information?”*

*Change the temperature to a temperature that you think will help the model to respond more accurately; higher temperature allows creativity while lower temperature encourages conservative answers and consistency*.

*For your information, here are previous system contents separated by ‘---’:*

## Appendix 4: Prompt for Complex Gene Pathways Analysis

Query: As a molecular biology expert, what is the relationship between gene “A” and gene “B” in the Non-small Cell Lung Cancer gene pathway? It’s possible that these two genes are connected by multiple intermediate genes. Please describe the relationship connecting each gene with one of the following: activation, inhibition, phosphorylation, and ubiquitination. Please also provide references if available. After the knowledge is extracted, please write the knowledge in the form of Cypher script ready to be used for visualization in Neo4j. Moreover, two genes may be connected by multiple distinct pathways; please generate your answer covering all the pathways between two genes. Make sure your answer is exactly correct.

Q: PI3K and AKT

A: PI3K activates PIP3, PIP3 activates AKT. Therefore, PI3K -[activates]-> PIP3 - [activates]-> AKT. The Cypher Script: (A: gene {name:”PI3K”}) -[:activates {source:”ChatGPT”}]->(B:gene {name:”PIP3”}) -[:activates {source:”ChatGPT”}]-> (C:gene {name: “AKT”})

Q: RAF and CCND1

A: RAF activates MEK, MEK activates ERK, ERK activates CCND1. Therefore, RAF-[activates]->MEK-[activates]->ERK-[activates]->CCND1. The Cypher Script is: (A: gene {name:”RAF”}) -[:activates {source:”ChatGPT”}]->(B:gene {name:”MEK”}) - [:activates {source:”ChatGPT”}]-> (C:gene {name: “ERK”}) -[:activates {source:”ChatGPT”}]-> (D:gene {name:”CCND1”})

Q: RET and RAF

A: RET activates RAS, RAS activates RAF. Therefore, RET-[activates]->RAS- [activates]->RAF. The Cypher script is: (A: gene {name:”RET”}) -[:activates {source:”ChatGPT”}]->(B:gene {name:”RAS”}) -[:activates {source:”ChatGPT”}]-> (C:gene {name: “RAF”})

Q: EGFR and ERK

A:

ChatGPT (GPT-4): EGFR activates RAS, RAS activates RAF, RAF activates MEK, MEK activates ERK. Therefore, EGFR-[activates]->RAS-[activates]->RAF-[activates]->MEK-[activates]->ERK. The Cypher Script is: (A: gene {name:”EGFR”}) -[:activates {source:”ChatGPT”}]->(B:gene {name:”RAS”}) -[:activates {source:”ChatGPT”}]-> (C:gene {name: “RAF”}) -[:activates {source:”ChatGPT”}]-> (D:gene {name:”MEK”}) - [:activates {source:”ChatGPT”}]-> (E:gene {name:”ERK”})

Query: Why do you think so?

ChatGPT (GPT-4): My response is based on well-established molecular biology knowledge and various research findings. The relationships I described are part of the canonical signaling pathways involving these genes, which have been widely studied in Non-small Cell Lung Cancer (NSCLC) and other cancers.

EGFR (Epidermal Growth Factor Receptor) is a receptor tyrosine kinase that activates the RAS-RAF-MEK-ERK signaling pathway upon activation by a ligand. This pathway is crucial in regulating cell proliferation, differentiation, and survival. In NSCLC, mutations in EGFR and other components of this pathway can lead to aberrant signaling and cancer progression.

Here are some references to support this information:

1. Lemmon, M. A., & Schlessinger, J. (2010). Cell signaling by receptor tyrosine kinases. Cell, 141(7), 1117-1134. doi:10.1016/j.cell.2010.06.011
2. Dhillon, A. S., Hagan, S., Rath, O., & Kolch, W. (2007). MAP kinase signaling pathways in cancer. Oncogene, 26(22), 3279-3290. doi:10.1038/sj.onc.1210421
3. Downward, J. (2003). Targeting RAS signaling pathways in cancer therapy. Nature Reviews Cancer, 3(1), 11-22. doi:10.1038/nrc969

These references provide evidence for the relationships I described between EGFR, RAS, RAF, MEK, and ERK, and their roles in NSCLC and other cancer types.

## Appendix 5: Prompt for Gene Pathways Reconstruction

Query: Can you rebuild the gene pathway of non-small cell lung cancer? Please generate the Cypher Script that can be used to reconstruct the graph in Neo4j as well.

ChatGPT (GPT-4): Reconstructing the gene pathway of non-small cell lung cancer (NSCLC) involves understanding the various genes and mutations involved in the disease. While I am unable to provide you with a comprehensive gene pathway, I can offer you an example Cypher script that demonstrates how to create a simplified graph in Neo4j. Please note that this example is not exhaustive, and you may need to consult the latest research for a complete representation of the gene pathway.

cypher

// Create genes

CREATE (EGFR:Gene {name: ‘EGFR’, description: ‘Epidermal growth factor receptor’})

CREATE (KRAS:Gene {name: ‘KRAS’, description: ‘Kirsten rat sarcoma viral oncogene’})

CREATE (ALK:Gene {name: ‘ALK’, description: ‘Anaplastic lymphoma kinase’})

CREATE (MET:Gene {name: ‘MET’, description: ‘MET proto-oncogene, receptor tyrosine kinase’})

CREATE (BRAF:Gene {name: ‘BRAF’, description: ‘B-Raf proto-oncogene, serine/threonine kinase’})

CREATE (ROS1:Gene {name: ‘ROS1’, description: ‘ROS proto-oncogene 1, receptor tyrosine kinase’})

CREATE (RET:Gene {name: ‘RET’, description: ‘Ret proto-oncogene’})

// Create NSCLC entity

CREATE (NSCLC:Cancer {name: ‘Non-small cell lung cancer’})

// Create relationships between genes and NSCLC

CREATE (EGFR)-[:INVOLVED_IN]->(NSCLC)

CREATE (KRAS)-[:INVOLVED_IN]->(NSCLC)

CREATE (ALK)-[:INVOLVED_IN]->(NSCLC)

CREATE (MET)-[:INVOLVED_IN]->(NSCLC)

CREATE (BRAF)-[:INVOLVED_IN]->(NSCLC)

CREATE (ROS1)-[:INVOLVED_IN]->(NSCLC)

CREATE (RET)-[:INVOLVED_IN]->(NSCLC)

This script creates nodes for some key genes associated with NSCLC and a node for the cancer itself. It then defines relationships between these genes and NSCLC. Please keep in mind that this example is for illustrative purposes and does not capture the full complexity of NSCLC’s gene pathway.

## Appendix 6: Least to Most Prompt for Pathway Reconstruction

Query: As a molecular biology and bioinformatics expert. Can you reconstruct the Non-small Cell Lung Cancer gene pathway? Make sure your answer has a good confidence. Please generate the Cypher Script that can be used to reconstruct the graph in Neo4j as well. The relation connecting each gene should be specified with one of the following words: activates, inhibits and phosphorylates. What subquestions should be solved to help you build a more complete gene pathway graph?

ChatGPT (GPT-4): To build a more complete gene pathway graph for NSCLC, you can consider solving the following subquestions:

1. Which additional genes are involved in the NSCLC pathway and their respective roles?
2. How do these genes interact with each other, considering different types of relationships (activates, inhibits, phosphorylates)?
3. How do different mutations in these genes contribute to the development of NSCLC and the patient’s response to treatment?
4. How do the different NSCLC subtypes (adenocarcinoma, squamous cell carcinoma, and large cell carcinoma) influence the gene pathway and interactions?
5. Are there any potential drug targets or biomarkers that can be identified within the NSCLC gene pathway?

Answering these subquestions can provide a more comprehensive understanding of the NSCLC gene pathway and may help in the development of targeted therapies for this cancer type.

Query: Inquiry: As a molecular biology and bioinformatics expert. Can you reconstruct the Non-small Cell Lung Cancer gene pathway?

Instruction: Make sure your answer has a good confidence. Please generate the Cypher Script that can be used to reconstruct the graph in Neo4j as well. The relation connecting each gene should be specified with one of the following words: activates, inhibits and phosphorylates.

Q: Which additional genes are involved in the NSCLC pathway and their respective roles?

ChatGPT (GPT-4): MET: This gene encodes the hepatocyte growth factor receptor and has been implicated in NSCLC progression. MET amplification or mutation can lead to increased cell proliferation, survival, and angiogenesis.

RET: This gene encodes the RET receptor tyrosine kinase, and its rearrangement has been associated with a subset of NSCLC cases. RET fusions lead to constitutive activation of downstream signaling pathways.

ERBB2 (HER2): This gene encodes a member of the epidermal growth factor receptor (EGFR) family. ERBB2 amplification and overexpression have been observed in a subset of NSCLC patients, mainly in adenocarcinoma.

NTRK1, NTRK2, NTRK3: These genes encode tropomyosin receptor kinases (TRKs), and their fusions have been identified in a small subset of NSCLC patients. TRK fusions lead to oncogenic activation and can drive tumor growth.

RICTOR: This gene is a component of the mammalian target of rapamycin complex 2 (mTORC2) and is involved in the regulation of cell growth, proliferation, and survival. RICTOR amplification has been observed in NSCLC and is associated with poor prognosis.

…

(Process truncated because of space issue)

ChatGPT (GPT-4): Below is the NSCLC gene pathway reconstruction in the form of a Cypher script that can be used to create the graph in Neo4j. This script includes the main genes and their interactions, using “activates”, “inhibits”, and “phosphorylates” relationships:

CREATE (MET:Gene {name: ‘MET’})

CREATE (RET:Gene {name: ‘RET’})

CREATE (ERBB2:Gene {name: ‘ERBB2’})

CREATE (NTRK1:Gene {name: ‘NTRK1’})

CREATE (NTRK2:Gene {name: ‘NTRK2’})

CREATE (NTRK3:Gene {name: ‘NTRK3’})

CREATE (RICTOR:Gene {name: ‘RICTOR’})

CREATE (PI3K:Gene {name: ‘PI3K’})

CREATE (AKT:Gene {name: ‘AKT’})

CREATE (mTOR:Gene {name: ‘mTOR’})

CREATE (RAS:Gene {name: ‘RAS’})

CREATE (RAF:Gene {name: ‘RAF’})

CREATE (MEK:Gene {name: ‘MEK’})

CREATE (ERK:Gene {name: ‘ERK’})

CREATE (PLCg:Gene {name: ‘PLC_γ_’})

CREATE (PKC:Gene {name: ‘PKC’})

CREATE

(MET)-[:ACTIVATES]->(PI3K),

(MET)-[:ACTIVATES]->(RAS),

(RET)-[:ACTIVATES]->(RAS),

(RET)-[:ACTIVATES]->(PI3K),

(ERBB2)-[:ACTIVATES]->(PI3K),

(ERBB2)-[:ACTIVATES]->(RAS),

(NTRK1)-[:ACTIVATES]->(PI3K),

(NTRK1)-[:ACTIVATES]->(RAS),

(NTRK1)-[:ACTIVATES]->(PLCg),

(NTRK2)-[:ACTIVATES]->(PI3K),

(NTRK2)-[:ACTIVATES]->(RAS),

(NTRK2)-[:ACTIVATES]->(PLCg),

(NTRK3)-[:ACTIVATES]->(PI3K),

(NTRK3)-[:ACTIVATES]->(RAS),

(NTRK3)-[:ACTIVATES]->(PLCg),

(RICTOR)-[:PHOSPHORYLATES]->(AKT),

(PI3K)-[:ACTIVATES]->(AKT),

(AKT)-[:ACTIVATES]->(mTOR),

(RAS)-[:ACTIVATES]->(RAF),

(RAF)-[:ACTIVATES]->(MEK),

(MEK)-[:ACTIVATES]->(ERK),

(PLCg)-[:ACTIVATES]->(PKC)

;

This script creates nodes for the genes involved in the NSCLC pathway and their interactions, focusing on the PI3K/AKT/mTOR, RAS/RAF/MEK/ERK, and PLC_γ_-PKC pathways. Please note that this is a simplified representation of the complex interactions occurring in the NSCLC pathway.

## Appendix 7: Tree of Thoughts Tables

**Appendix Table 1.**
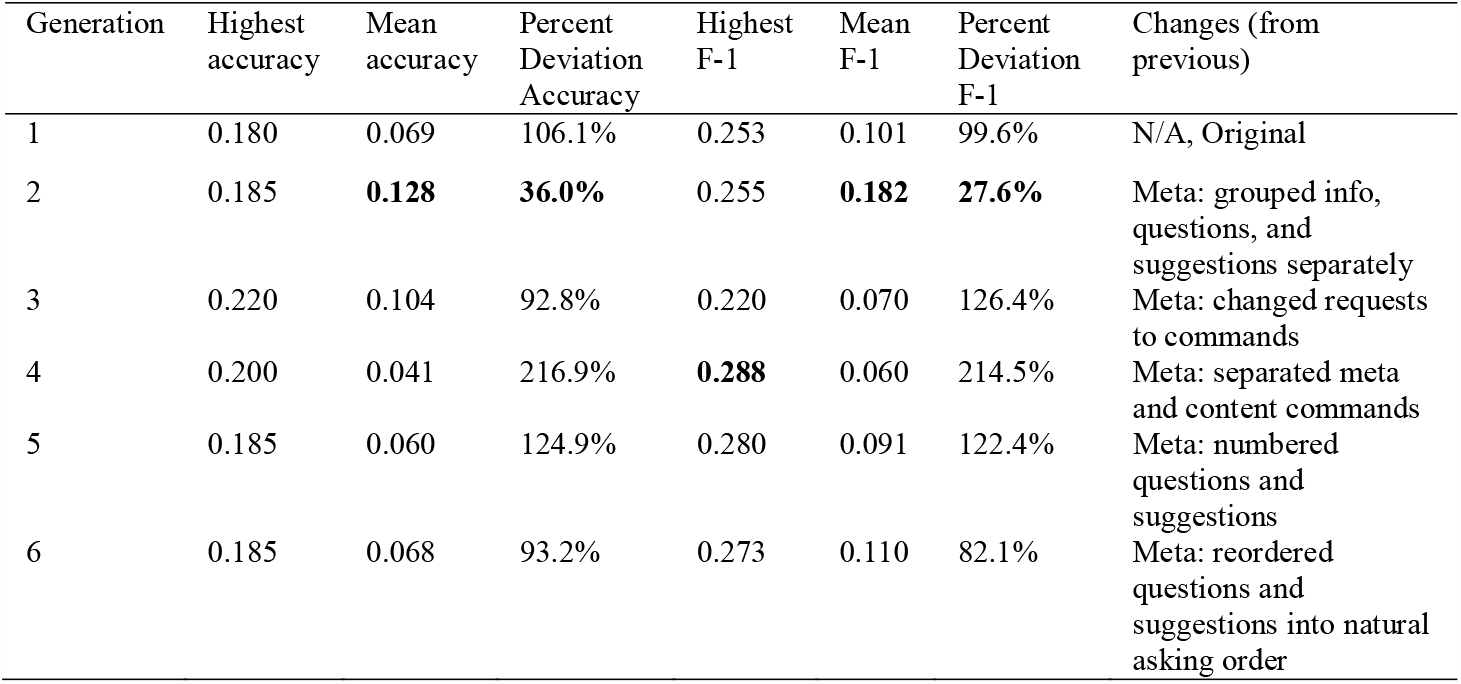
Evolution of Meta-Prompt Refinement Across Different Generations. The table presents performance metrics such as highest accuracy, mean accuracy, percent deviation of accuracy, highest F-1, mean F-1, and percent deviation of F-1. The bold fonts indicate the best performance of each column. Each row represents distinct iterations, highlighting specific changes made to the meta-prompt in relation to prior iterations and their corresponding performance impact.

**Appendix Table 2.**
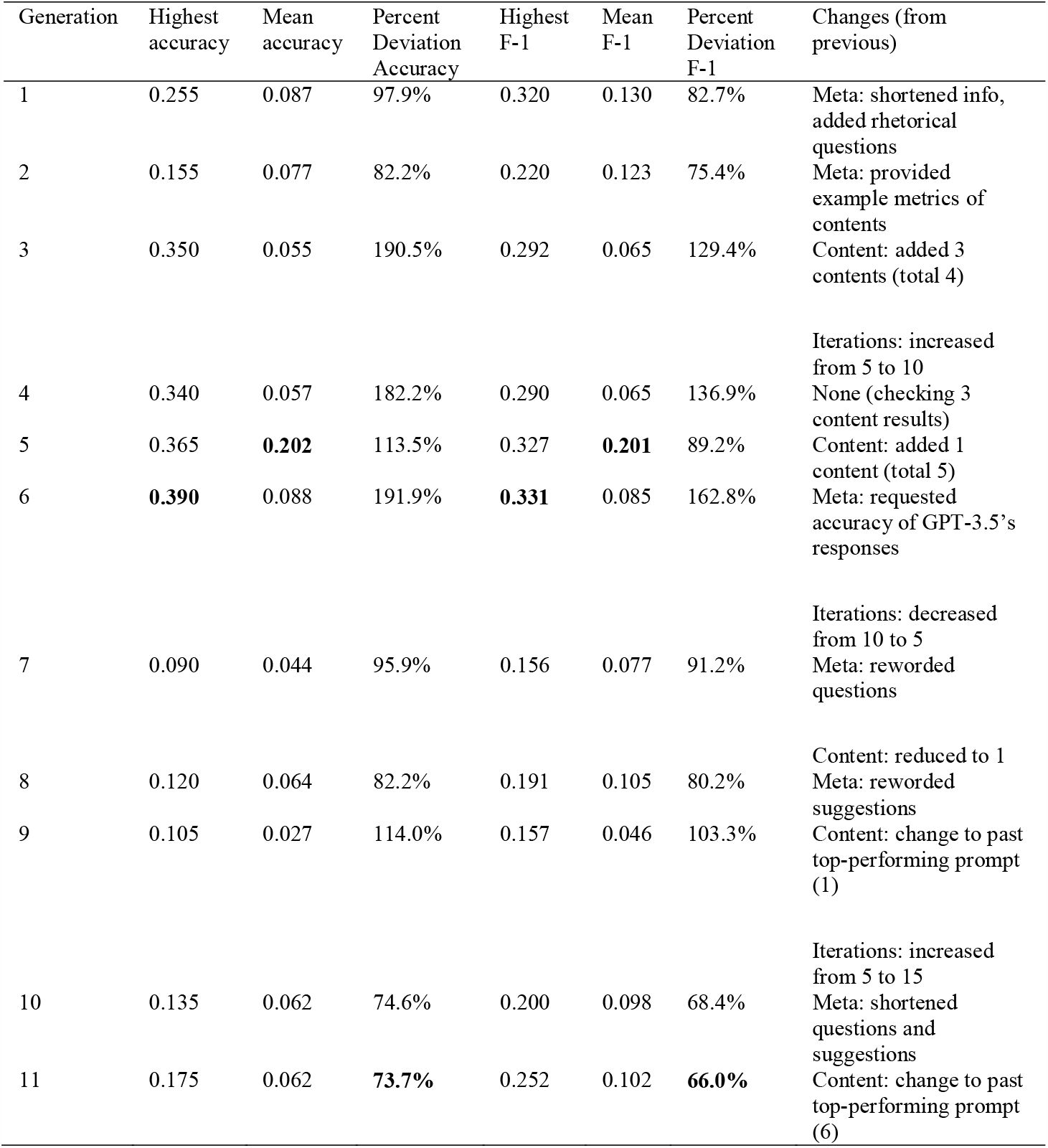
Influence of System Content Adjustments on Model Performance Across Generations. The table enumerates the performance metrics for various generational iterations, including highest accuracy, mean accuracy, percent deviation of accuracy, highest F-1, mean F-1, and percent deviation of F-1. The bold fonts indicate the best performance of each column. Each iteration reflects specific modifications to the meta-prompt and system content, underscoring the relationship between content adjustments and performance shifts. The entries encapsulate content addition or modifications, prompt strategy variations, and iteration count alterations, providing insights into their respective impacts on the model’s efficacy.

| Generation | Highest accuracy | Mean accuracy | Percent Deviation Accuracy | Highest F-1 | Mean F-1 | Percent Deviation F-1 | Changes (from previous) |
|---|---|---|---|---|---|---|---|
| 1 | 0.255 | 0.087 | 97.9% | 0.320 | 0.130 | 82.7% | Meta: shortened info, added rhetorical questions |
| 2 | 0.155 | 0.077 | 82.2% | 0.220 | 0.123 | 75.4% | Meta: provided example metrics of contents |
| 3 | 0.350 | 0.055 | 190.5% | 0.292 | 0.065 | 129.4% | Content: added 3 contents (total 4) |
| 4 | 0.340 | 0.057 | 182.2% | 0.290 | 0.065 | 136.9% | Iterations: increased from 5 to 10<br>None (checking 3 content results) |
| 5 | 0.365 | **0.202** | 113.5% | 0.327 | **0.201** | 89.2% | Content: added 1 content (total 5) |
| 6 | **0.390** | 0.088 | 191.9% | **0.331** | 0.085 | 162.8% | Meta: requested accuracy of GPT-3.5’s responses |
| 7 | 0.090 | 0.044 | 95.9% | 0.156 | 0.077 | 91.2% | Iterations: decreased from 10 to 5<br>Meta: reworded questions |
| 8 | 0.120 | 0.064 | 82.2% | 0.191 | 0.105 | 80.2% | Content: reduced to 1<br>Meta: reworded suggestions |
| 9 | 0.105 | 0.027 | 114.0% | 0.157 | 0.046 | 103.3% | Content: change to past top-performing prompt (1) |
| 10 | 0.135 | 0.062 | 74.6% | 0.200 | 0.098 | 68.4% | Iterations: increased from 5 to 15<br>Meta: shortened questions and suggestions |
| 11 | 0.175 | 0.062 | **73.7%** | 0.252 | 0.102 | **66.0%** | Content: change to past top-performing prompt (6) |

**Appendix Table 3.**
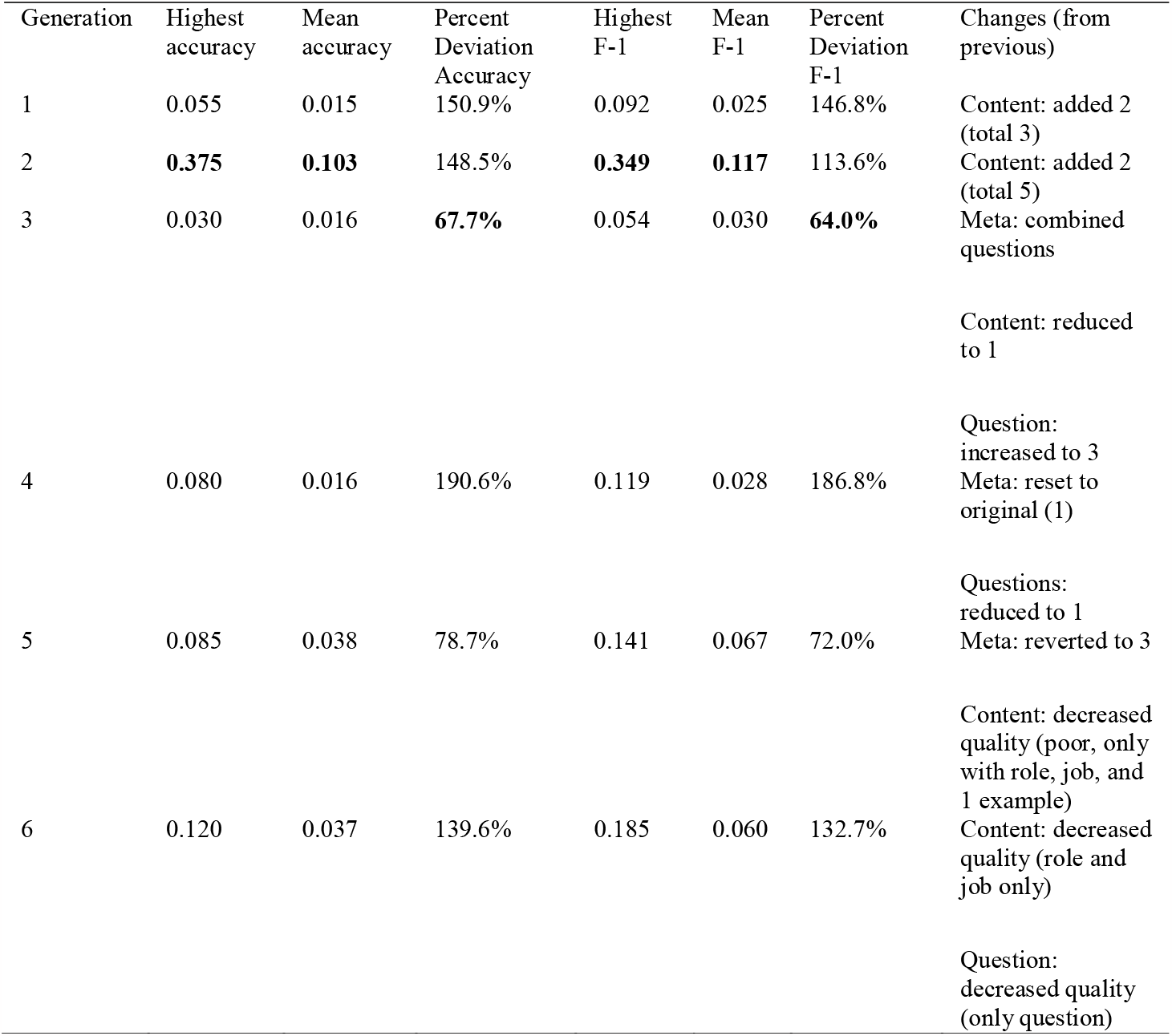
Comparative Analysis of the Impact of User Question Parameter on GPT-3.5’s Response Accuracy and F-1 Score. This table systematically summarizes GPT-3.5’s performance across different generations, emphasizing the highest and mean accuracy, percent deviation in accuracy, highest and mean F-1 scores, percent deviation in F-1, and corresponding content changes. The bold fonts indicate the best performance of each column. Notably, the data reinforces the observation that the quality of user questions plays a pivotal role in influencing the knowledge-mining efficacy of GPT-3.5.

## Notes

### Competing Interest Statement

The authors have declared no competing interest.

